# Temperature robustness of the timing network within songbird premotor nucleus HVC

**DOI:** 10.1101/2025.03.06.641874

**Authors:** Aayush Khare, Derek Sederman, Dezhe Z. Jin

**Affiliations:** Department of Physics and Huck Institute for Life Sciences, Pennsylvania State University, University Park, PA 16802, USA

## Abstract

Many neuronal processes are temperature-sensitive. Cooling by 10 °C typically slows ion channel dynamics by more than a factor of two (Q_10_ > 2). Nevertheless, behaviors can remain robust despite variations in brain temperature. For instance, cooling the premotor nucleus HVC in zebra finches by 10 °C slows song production by only a factor of Q_10_ ∼ 1.3. Here we examine the temperature robustness of the synaptic chain network within HVC. Burst spike propagation along such a chain network is postulated to control the tempo of the song. We show that the dynamics of this network are resilient to cooling and that the slowing of burst propagation exhibits a Q_10_ similar to that observed for the song. We identify two key factors underlying this robustness: the reliance on axonal delays, which are more resistant to temperature changes than ion channels, and enhanced synaptic efficacy at lower temperatures. We propose that these mechanisms represent general principles by which neural circuits maintain functional stability despite temperature fluctuations in the brain.

**Significance Statement:** Many animal behaviors remain robust despite temperature fluctuations in the brain. By studying timing circuits in songbirds, we identify key circuit elements that contribute to this resilience, including axonal delays and synaptic integration. Our work highlights how these mechanisms interact to maintain stable neuronal dynamics in response to temperature changes.

## Introduction

Temperature affects most neuronal processes. Cooling decreases the activation and deactivation rates of ion channels that govern neuron dynamics (Schauf, 1973; Schwarz and Eikhof, 1987; Lee et al., 2005), slows down spike propagation along the axon (Huxley, 1959; Swadlow et al., 1981; Janssen, 1992; Moran and Melani, 2001), and reduces both synaptic and ion channel conductance (Hodgkin et al., 1952; Sterratt, 2015). In cold-blooded animals, significant fluctuations in ambient temperature can challenge the neural circuits responsible for essential behaviors (Tang et al., 2010). In contrast, warm-blooded animals, such as mammals and birds, mitigate this challenge by maintaining a constant body temperature (Tan and Knight, 2018). However, brain temperature in these animals can still fluctuate by up to 4 °C (Moser et al., 1993; Kiyatkin, 2007; Kiyatkin, 2010; Aronov and Fee, 2012), leading to detectable changes in both behavior (Aronov and Fee, 2012) and neuronal dynamics (Petersen et al., 2022). Focal cooling of specific brain areas has also been used to establish causal links between brain regions and behaviors (Long and Fee, 2008; Aronov et al., 2011; Aronov and Fee, 2011; Goldin et al., 2013; Long et al., 2016; Zhang et al., 2017; Hamaguchi et al., 2016; Banerjee et al., 2019; Banerjee et al., 2021; Petersen et al., 2022).

The production of song in songbirds is controlled by the premotor nucleus HVC (proper name) and its downstream target, the robust nucleus of the arcopallium (RA) (Nottebohm et al., 1976) (Fig. 1a). Neurons in these nuclei exhibit precise spike timing that corresponds with the timing of song features. In zebra finches, HVC neurons that project to RA (HVC_RA_ neurons) burst exactly once at precise moments during a song motif (Hahnloser et al., 2002). Furthermore, the bursts of the HVC_RA_ neuron population form a continuous sequence that spans the entire song motif without gaps (Long et al., 2010; Lynch et al., 2016; Picardo et al., 2016; Egger et al., 2020).

**Figure 1:**
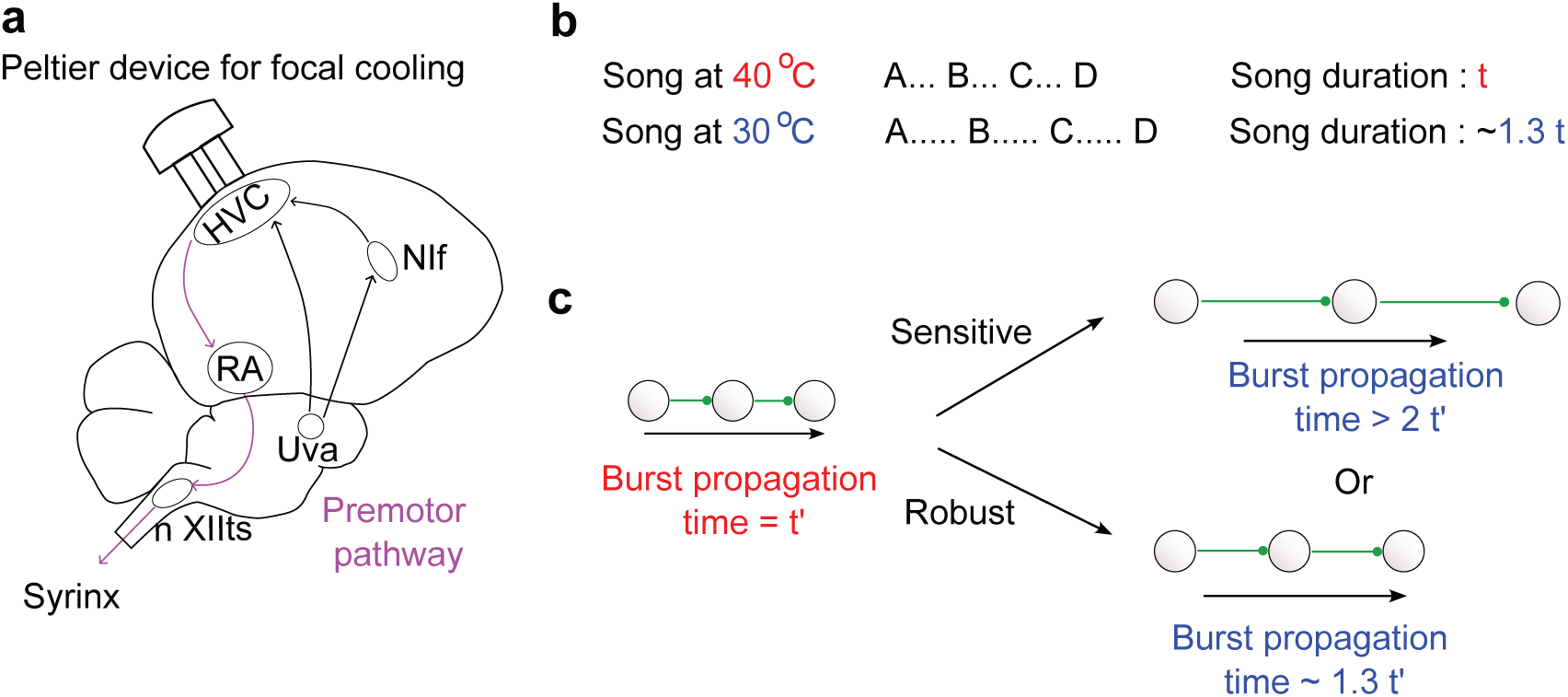
Temperature robustness of song tempo. (a) Song production is controlled by a feedforward pathway consisting of HVC (proper name), RA (the robust nucleus of the arcopallium), and nXIIts (the tracheosyringeal nucleus of the XII cranial nerve). Additionally, a feedback pathway involves Uva (the nucleus uvaeformis) and NIf (the nucleus interface). Cooling HVC with a Peltier device slows down the song. (b) Cooling HVC from its normal temperature (40 °C) by 10 °C increases the song duration by a factor of 1.3 (Q_10_ ∼ 1.3). (c) In our model, song tempo is regulated by a synaptic chain network of HVC_RA_ neurons. The central question of this study is whether this network is sensitive to or robust against temperature changes in HVC.

Cooling HVC by 10 °C slows the song by a factor of 1.3 (Fig. 1b). In contrast, cooling RA has minimal effect on song tempo (Long and Fee, 2008). These findings support the hypothesis that the ultra-sparse and precisely timed bursts of HVC_RA_ neurons are generated intrinsically within the HVC via a synaptic chain network (Jin et al., 2007; Long et al., 2010; Egger et al., 2020; Tupikov and Jin, 2021). HVC_RA_ neurons form a unidirectional synaptic chain network, where burst activity initiated at the start of the chain propagates along the network’s connections (Fig. 1c). Axonal delays between HVC_RA_ neurons, ranging from 1 to 7.5 ms, promote the formation of a polychronous network that uniformly distributes the burst times of HVC_RA_ neurons across the song motif (Egger et al., 2020; Tupikov and Jin, 2021).

This hypothesis was challenged based on a quantitative argument regarding the effects of HVC cooling. (Hamaguchi et al., 2016). The temperature sensitivity of a biological process is commonly quantified using a Q_10_ value, defined as the ratio of the rate of the process when the temperature changes by 10 °C (Huxley, 1959). A Q_10_ factor close to 1 indicates high temperature robustness. Neuronal processes, such as the activation and deactivation rates of ion channels, typically slow down with cooling, exhibiting Q_10_ values greater than 2 (Schauf, 1973; Schwarz and Eikhof, 1987; Lee et al., 2005). This raises a critical question: Why does birdsong tempo slow with a Q_10_ value of approximately 1.3 instead of Q_10_ > 2, as one might intuitively expect? (Hamaguchi et al., 2016) (Fig. 1c). Based on this apparent discrepancy, it was proposed that the HVC may be only one component of a distributed network involving multiple brain areas that form a loop to generate the moment-to-moment sequential activations of HVC_RA_ neurons (Hamaguchi et al., 2016).

In this study, we use computational modeling to show that the dynamics of the synaptic chain network localized within the HVC are as robust as the song itself against HVC cooling. We identify two key mechanisms. First, spike propagation along axons is robust to temperature changes, with Q_10_ ∼ 1.25 (Swadlow et al., 1981). Thus, the reliance on substantial axonal delays between HVC_RA_ neurons (Egger et al., 2020; Tupikov and Jin, 2021) contributes to the temperature robustness of the synaptic chain network. Second, cooling enhances the efficacy of synaptic transmission. The decay dynamics of synaptic inputs are highly temperature-sensitive, slowing down with Q_10_ > 2 when cooled (Gardner, 1980; Hestrin et al., 1990). This prolongs the duration of synaptic integration (Andersen and Moser, 1995). Coupled with the rise of the resting membrane potential and a reduction in leak conductance, the extended synaptic integration broadens excitatory postsynaptic potentials (EPSPs) despite the reduction in synaptic conductance (Volgushev et al., 2000). Hence, cooling enhances synaptic efficacy between HVC_RA_ neurons. It has been shown that excitatory inputs from the nucleus interface (NIf) elevate the membrane potentials of HVC_RA_ neurons during singing (Otchy et al., 2015). Cooling further increases the efficacy of these inputs. These factors collectively counteract the slowing of ion channel dynamics in HVC_RA_ neurons.

Our findings provide strong evidence that HVC is the primary site responsible for generating timing signals in birdsong. The two key factors we identify may also play a crucial role in maintaining the robustness of behaviors in other animals, such as spatial learning in rodents, despite temperature changes in the brain (Andersen and Moser, 1995).

## Results

The normal temperature of the zebra finch HVC is 40 °C (Long and Fee, 2008; Aronov and Fee, 2012). In this study, we investigate the changes in HVC dynamics when cooled to 30 °C.

### Synaptic chain network model

In our model, the network responsible for generating precisely timed burst spikes consists of 2000 HVC_RA_ neurons organized into a feedforward synaptic chain, similar to previous models (Jin et al., 2007; Long et al., 2010; Egger et al., 2020; Tupikov and Jin, 2021). Additionally, we include 550 HVC_INT_ neurons to incorporate feedback inhibition into the network (Fig. 2a). Axonal delays between HVC_RA_ neurons follow a log-normal distribution, ranging from 1.2 to 5.9 ms (5th to 95th percentile, Supplementary Fig. S1), consistent with experimental observations (Egger et al., 2020). Similarly, axonal delays from HVC_RA_ to HVC_INT_ neurons and from HVC_INT_ to HVC_RA_ neurons also follow log-normal distributions, ranging from 0.8 to 2.7 ms and 0.5 to 2.0 ms (5th to 95th percentile), respectively (Supplementary Fig. S1).

**Figure 2:**
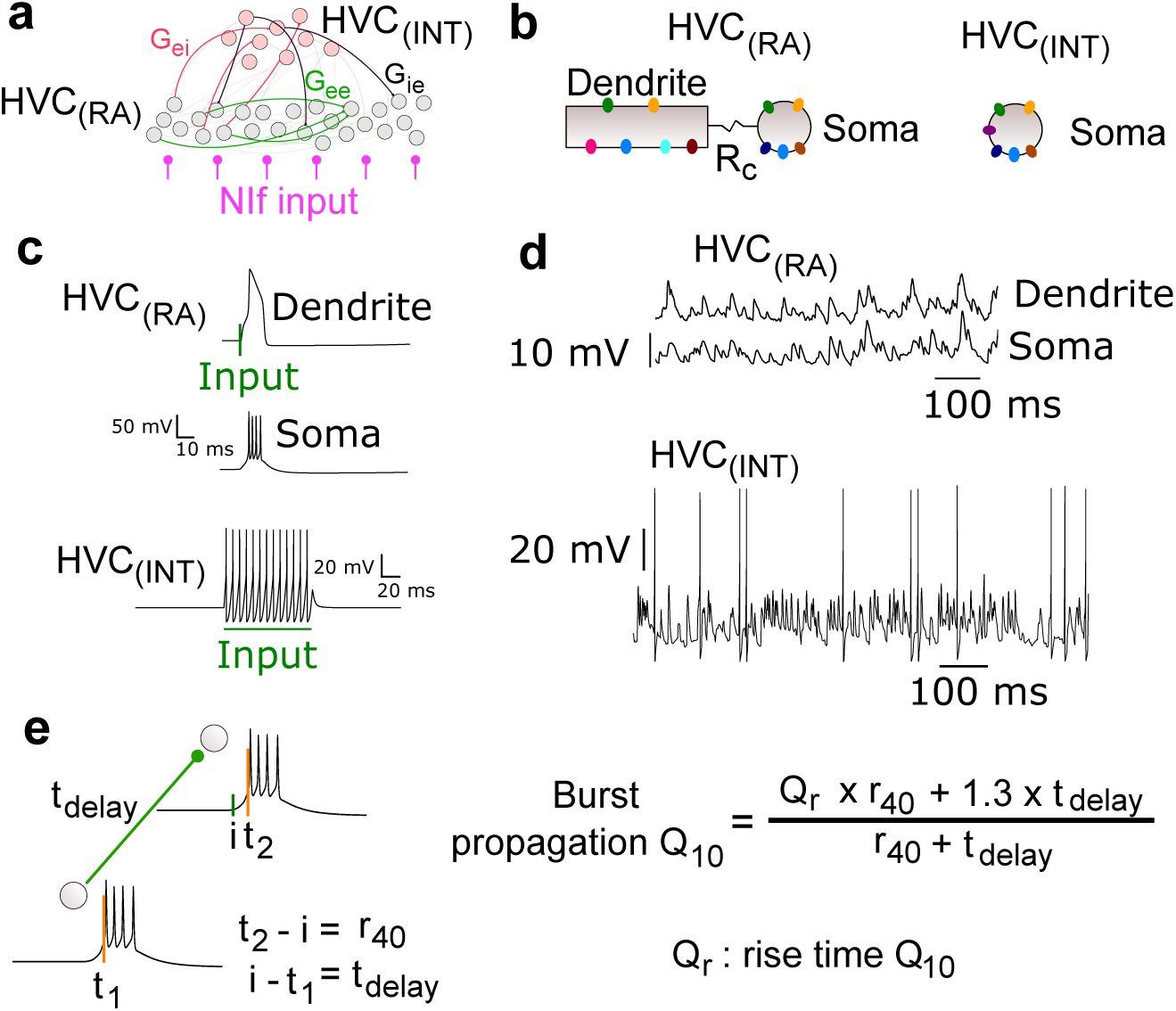
The synaptic chain network model. **a**. The HVC_RA_ neurons form a polychronous synaptic chain network. The HVC_INT_ neurons receive excitatory input from HVC_RA_ neurons and provide feedback inhibition to HVC_RA_ neurons. **b**. The HVC_RA_ neuron model consists of a dendritic compartment and a somatic compartment that are ohmically coupled. The HVC_INT_ neuron is modeled as a single-compartment neuron. **c**. Strong excitatory input to the HVC_RA_ dendrite induces a calcium spike, which drives a stereotypical burst of action potentials in the soma. Current input to the HVC_INT_ neuron results in high-frequency spiking activity. **d**. Noise induces subthreshold fluctuations in both compartments of HVC_RA_ neurons and spontaneous spike activity in HVC_INT_ neurons. **e**. Burst propagation between two HVC_RA_ neurons involves two timescales: axonal delay and subthreshold integration.

Despite these distributed axonal delays, post-synaptic HVC_RA_ neurons receive synchronous inputs, consistent with the polychronous principle (Izhikevich, 2006; Egger et al., 2020; Tupikov and Jin, 2021). Such a polychronous chain network naturally emerges through a self-organizing process involving synaptic plasticity and spontaneous activity (Tupikov and Jin, 2021). In this study, we use a simplified approach to wire the polychronous chain network, ensuring nearly synchronous inputs and generating smooth burst sequences of HVC_RA_ neurons (Methods, Supplementary Fig. S1a). Additionally, we wire HVC_INT_ neurons using a method that creates gaps in HVC_INT_ neuron activity before the burst times of HVC_RA_ neurons (Methods, Supplementary Fig. S1b, c), in accordance with experimental observations of HVC_RA_ and HVC_INT_ neuron firing patterns during singing (Kosche et al., 2015).

The HVC_RA_ neuron model consists of a dendritic compartment and a somatic compartment (Fig. 2b). Both compartments include a leak conductance. Additionally, the dendritic compartment contains calcium conductance and calcium-activated potassium conductance, enabling the generation of a calcium spike in response to strong input (Jin et al., 2007; Long et al., 2010). The somatic compartment contains both sodium and delayed-rectifier potassium conductances, which facilitate the generation of sodium spikes. When driven by a dendritic spike, the soma produces stereotypical bursts of 4–5 sodium spikes (Fig. 2c).

The HVC_INT_ neuron is modeled as a single-compartment neuron (Fig. 2b) (Jin et al., 2007; Long et al., 2010). In addition to the leak, sodium, and delayed-rectifier potassium currents, it includes a high-threshold potassium current that enables spiking at high frequencies (Fig. 2c).

Noise is added to the neurons. At 40 °C, the membrane potentials of HVC_RA_ neurons fluctuate with a standard deviation of around 3 mV (Fig. 2d); and HVC_INT_ neurons spontaneously spike at around 10 Hz (Fig. 2d). Both values are derived from empirical observations (Mooney, 2000; Rauske et al., 2003; Long et al., 2010; Hamaguchi et al., 2016; Vallentin and Long, 2015; Kozhevnikov and Fee, 2007; Hozhabri et al., 2025).

### Effects of temperature on the model parameters

The effects of temperature on neuronal processes can be categorized into three groups. The first category includes processes that depend on protein conformational changes, such as the opening and closing dynamics of ion channels and the closing dynamics of synaptic receptors. These processes are highly temperature-sensitive, with cooling significantly slowing their dynamics (Q_10_ > 2) (Huxley, 1959; Schauf, 1973; Schwarz and Eikhof, 1987). In our model, we set Q_10_ = 3 for the rates of ion channel dynamics and the decay time of synaptic conductance.

The second category includes processes involving ion diffusion driven by voltage gradients in electrolytes (Armstrong and Hille, 1998). Ion channel conductance, synaptic conductance, and the conduction of spikes in unmyelinated axons belong to this category. These processes are more temperature-robust, with Q_10_ factors of approximately 1.3 (Hodgkin et al., 1952; Sterratt, 2015). In our model, cooling reduces all conductances and increases axonal delays with Q_10_ = 1.3.

Finally, the reversal potentials for ion channels are directly proportional to the absolute temperature (Dayan and Abbott, 2005). Therefore, cooling from 40 °C to 30 °C results in scaling the reversal potentials by a factor of (273+30)/(273+40) = 0.97.

The scaling of all cellular parameters from 40 °C to 30 °C is summarized in Table 1.

**Table 1:**
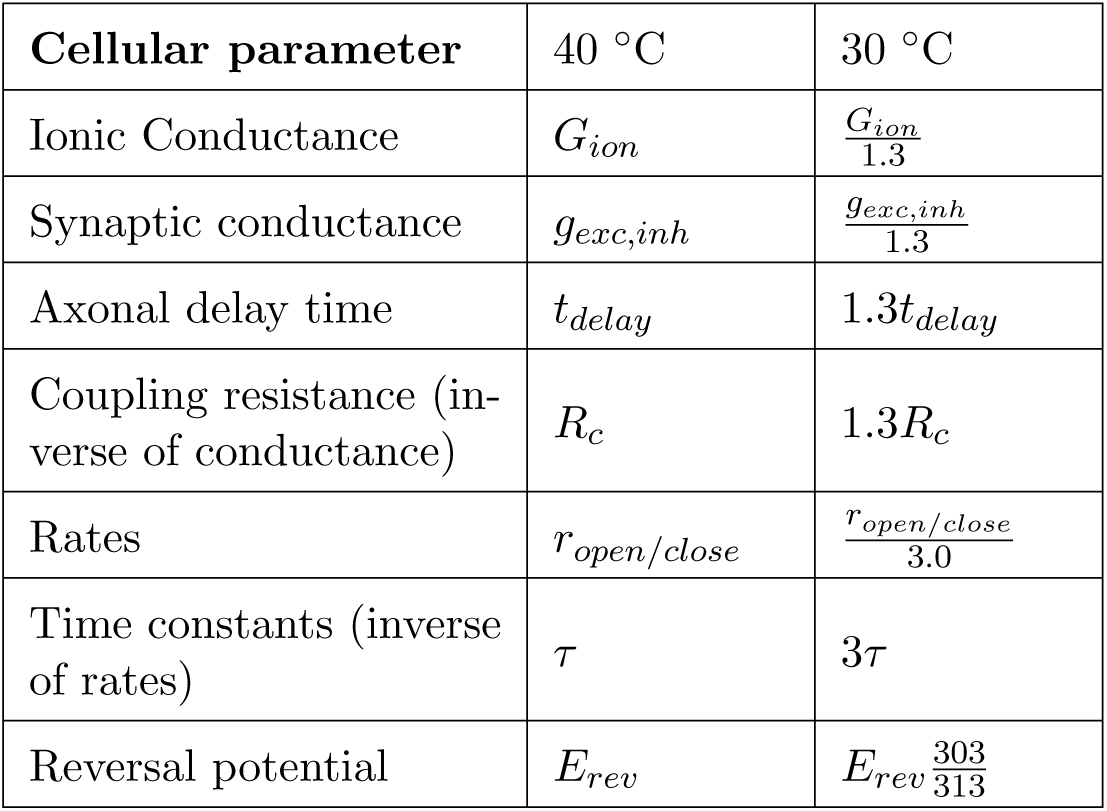
Changes of model parameters when temperature changed from 40 °C to 30 °C.

| Cellular parameter | 40 °C | 30 °C |
| --- | --- | --- |
| Ionic Conductance | $G_{ion}$ | $\frac{G_{ion}}{1.3}$ |
| Synaptic conductance | $g_{exc,inh}$ | $\frac{g_{exc,inh}}{1.3}$ |
| Axonal delay time | $t_{delay}$ | $1.3t_{delay}$ |
| Coupling resistance (inverse of conductance) | $R_c$ | $1.3R_c$ |
| Rates | $r_{open/close}$ | $\frac{r_{open/close}}{3.0}$ |
| Time constants (inverse of rates) | $\tau$ | $3\tau$ |
| Reversal potential | $E_{rev}$ | $E_{rev} \frac{303}{313}$ |

### Burst propagation Q_10_

At 40 °C, the time from the presynaptic HVC_RA_ burst to the postsynaptic HVC_RA_ burst consists of the axonal delay *t*_delay_ and the rise time *r*_40_ of the postsynaptic neuron’s somatic membrane potential (Fig. 2e). Upon cooling to 30 °C, this burst propagation time becomes

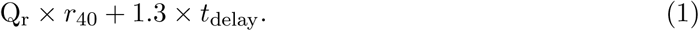

Here Q_r_ = *r*_30_*/r*_40_, where *r*_30_ is the rise time of the soma’s membrane potential at 30 °C. The Q_10_ factor for axonal delay time is 1.3 (Swadlow et al., 1981). Therefore, the Q_10_ factor of burst propagation time is

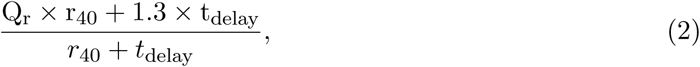

which is a weighted average of Q_r_ and 1.3. If *t*_delay_ ≫ *r*_40_, the Q_10_ factor is closer to 1.3. Conversely, if *t*_delay_ ≪ *r*_40_, the Q_10_ factor is closer to Q_r_. Experiments have shown that *t*_delay_ is in the range of 1 - 7.5 ms (Egger et al., 2020), while *r*_40_ is in the range of 5 - 10 ms (Long et al., 2010). These values suggest that axonal delay and the rise time of the soma’s membrane potential contribute to the burst propagation Q_10_ factor approximately equally.

### Synaptic response of an HVC_RA_ neuron

To understand Q_r_, we examined the response of an HVC_RA_ neuron when an excitatory synaptic input with conductance *G_e_* was delivered to its dendrite (Fig. 3a). The rise time, *r_T_*, was defined as the duration from the input onset to the peak of the somatic membrane potential if the response was subthreshold, or to the time of crossing -20 mV if the response was a burst of spikes (Fig. 3a). The factor Q_r_ depends on *G_e_*, and three distinct regions can be identified (Fig. 3b).

- For *G_e_ <* 0.29 mS/cm^2^ (region I), the neuron did not spike at either 40 °C or 30 °C. In this region, as *G_e_*increased, Q_r_ rose from 1.73 to 3.45.
- For 0.29 mS/cm^2^ *< G_e_ <* 0.53 mS/cm^2^ (region II), the neuron did not spike at 40 °C but did spike at 30 °C. This effect arises from two factors. First, cooling reduces the leak conductance with Q_10_ = 1.3, and the resting membrane potential is slightly elevated due to a higher leak reversal potential, making the neuron more excitable (Fig. 3c). Second, although the synaptic conductance scales down by 1.3, the synaptic decay time constant increases by a factor of Q_10_ = 3. This results in a broadened and enhanced EPSP at 30 °C. Consequently, the membrane potential reaches higher peak values, leading to a subthreshold-to-superthreshold transition at 30 °C but not at 40 °C. This transition occurs at the left edge of region II. As *G_e_*increases, *r*_30_ decreases faster than *r*_40_, causing Q_r_ to decline from 3.45 to 0.48.
- For *G_e_ >* 0.53 mS/cm^2^ (region III), the subthreshold-to-superthreshold transition also occurs at 40 °C. As *G_e_* increases, Q_r_ rises from 0.48 and eventually plateaus at 1.42.

**Figure 3:**
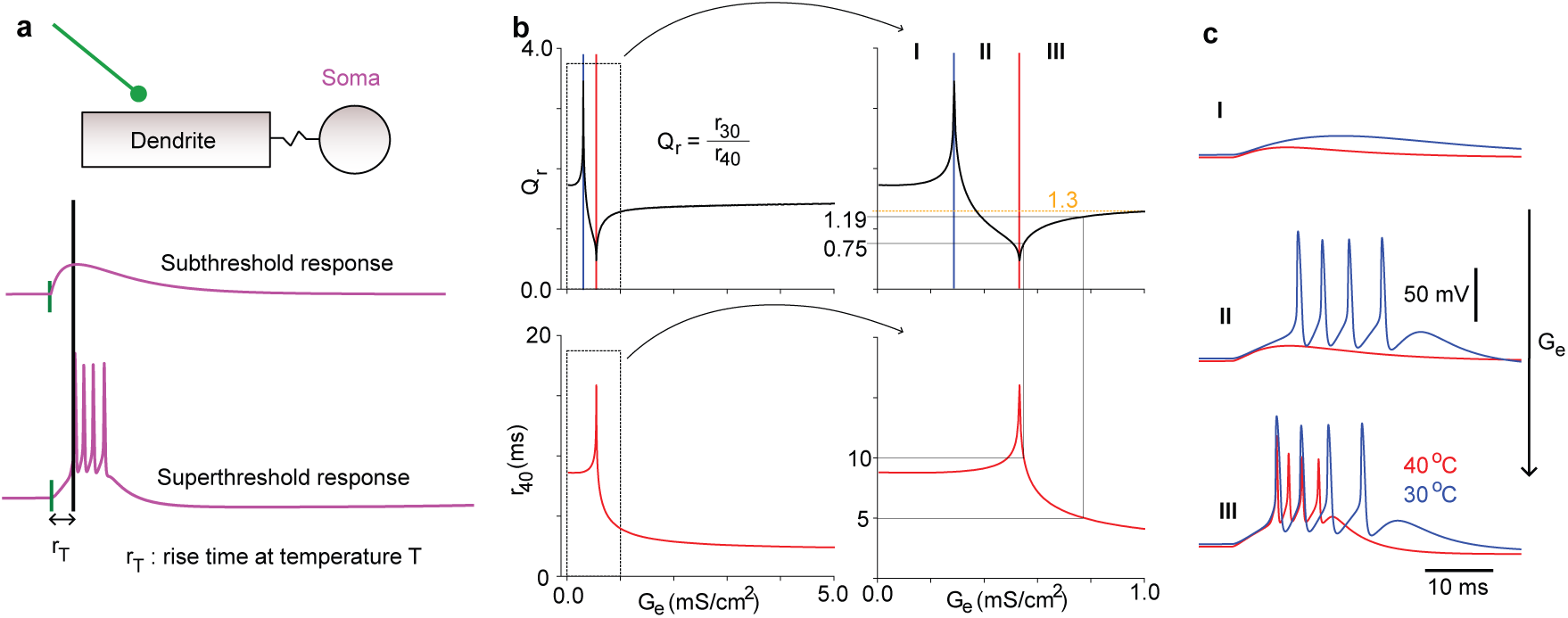
The synaptic response of an HVC_RA_ neuron. (a) A single excitatory synaptic input is applied at the dendrite of an HVC_RA_ neuron, and the rise time (*r_T_*) of the soma at a given temperature *T* is tracked. (b) The rise time *r*_40_ and its Q_10_ factor (Q_r_) as a function of the excitatory input strength *G_e_*. Three regions are identified based on the neuronal response at two different temperatures. The right panel shows a zoomed-in section of the excitation regime, highlighting a narrow segment in region III where the rise time falls within 5–10 ms. (c) Example somatic membrane potential traces corresponding to the three regions identified in the analysis.

Example traces of somatic membrane potentials across the three regions are shown in Fig. 3c.

Intracellular recordings in singing zebra finches indicate that the somatic membrane potential of HVC_RA_ neurons rises from baseline to spiking threshold within 5 - 10 ms (Long et al., 2010; Hamaguchi et al., 2016; Egger et al., 2020). In region III of our model, increasing *G_e_* decreases *r*_40_ (Fig. 3b). We find that *r*_40_ = 5 ms corresponds to Q_r_ = 1.19, while *r*_40_ = 10 ms corresponds to Q_r_ = 0.75. Thus, when *r*_40_ aligns with the experimental data, Q_r_ remains below 1.3 in our model.

These results can be understood through a simple mathematical analysis. In our model, the synaptic conductance follows a kick-and-decay dynamic (Methods). When a spike arrives, the conductance increases by *G_e_* and then decays according to the synaptic decay time constant *τ_s_*.

The time course of the synaptic conductance is therefore given by

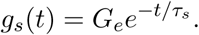

The subthreshold membrane potential of the dendrite can be approximated using leaky integration:

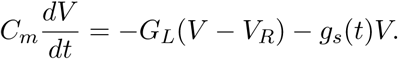

Here, *C_m_* is the membrane capacitance, *G_L_* is the leak conductance, and *V_R_* is the resting membrane potential. The EPSP peaks at *t* = *t_m_* with *V* = *V_m_*. At this peak,

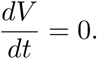

Thus, we obtain

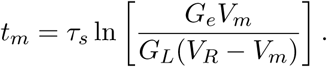

This equation shows that the synaptic decay time constant *τ_s_* is the primary factor determining *t_m_*. Therefore, the Q_10_ of *t_m_* should be approximately equal to that of *τ_s_*, which is 3 in our case. The temperature dependence of *G_e_* and *G_L_* cancels out. Thus, in the subthreshold regime (region I), the rise time *r_T_* is highly sensitive to temperature.

In this subthreshold regime, cooling increases the peak value *V_m_* of the EPSP. This occurs because the increase in the synaptic decay time constant, *τ_s_*, with a Q_10_ value of 3, outweighs the decrease in synaptic strength, *G_e_*, which has a Q_10_ value of 1.3. If the membrane potential is held constant, as in a voltage-clamp experiment, the charge transfer from the synapse is proportional to

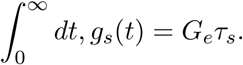

When cooled to 30 °C, this quantity increases by a factor of 3*/*1.3 = 2.3. Analyzing the case where the membrane potential *V* (*t*) is not clamped is more complex, but it is still possible to show mathematically that cooling increases *V_m_*for the same reason (see Methods).

The enhanced EPSP upon cooling enables the neuron to generate dendritic calcium spikes and somatic bursts at 30 °C but not at 40 °C when *G_e_* is further increased (region II), as shown in Fig. 3c. This dendritic spike contributes to reducing the rise time of the neuron at 30 °C, explaining why Q_r_ decreases in region II, as illustrated in Fig. 3b.

With sufficiently large *G_e_*, the neuron responds with a dendritic spike and a somatic burst at 40 °C as well (region III). Notably, there remains a range of *G_e_* where Q_r_ is less than 1. As *G_e_* increases further, Q_r_ rises but plateaus slightly above 1.4. In this regime, a dendritic spike is rapidly generated, which in turn drives a somatic burst of spikes through the ohmic coupling *R_c_* between the two compartments. Consequently, Q_r_ is largely determined by the Q_10_ of *R_c_* and the leak conductance *G_L_*, both of which have Q_10_ = 1.3. The initiation of calcium and sodium spikes is fast but has Q_10_ = 3, contributing to Q_r_ plateauing at a value slightly greater than 1.3.

### Effect of NIf inputs on the HVC_RA_ synaptic response

NIf provides major excitatory inputs to HVC (Lewandowski et al., 2013). Experiments involving the transient inactivation of NIf in zebra finches have demonstrated that NIf excitatory drive is essential for singing (Otchy et al., 2015). Interestingly, zebra finches can recover from NIf lesions, suggesting that the HVC network can homeostatically enhance connection strengths between HVC_RA_ neurons to compensate for the loss of NIf excitatory drive (Otchy et al., 2015). To investigate the impact of NIf excitatory drive on temperature robustness in our model, we repeated the study of the synaptic response in regime 3 (Fig. 3c) of an HVC_RA_ neuron, this time incorporating additional excitatory inputs to the dendritic compartment from NIf (Fig. 4a). The NIf input was modeled as random spikes generated by a Poisson process with a frequency of 1500 Hz. At 40 °C, this input elevated the resting membrane potential from -80 mV to -76.4 mV (Fig. 4b).

**Figure 4:**
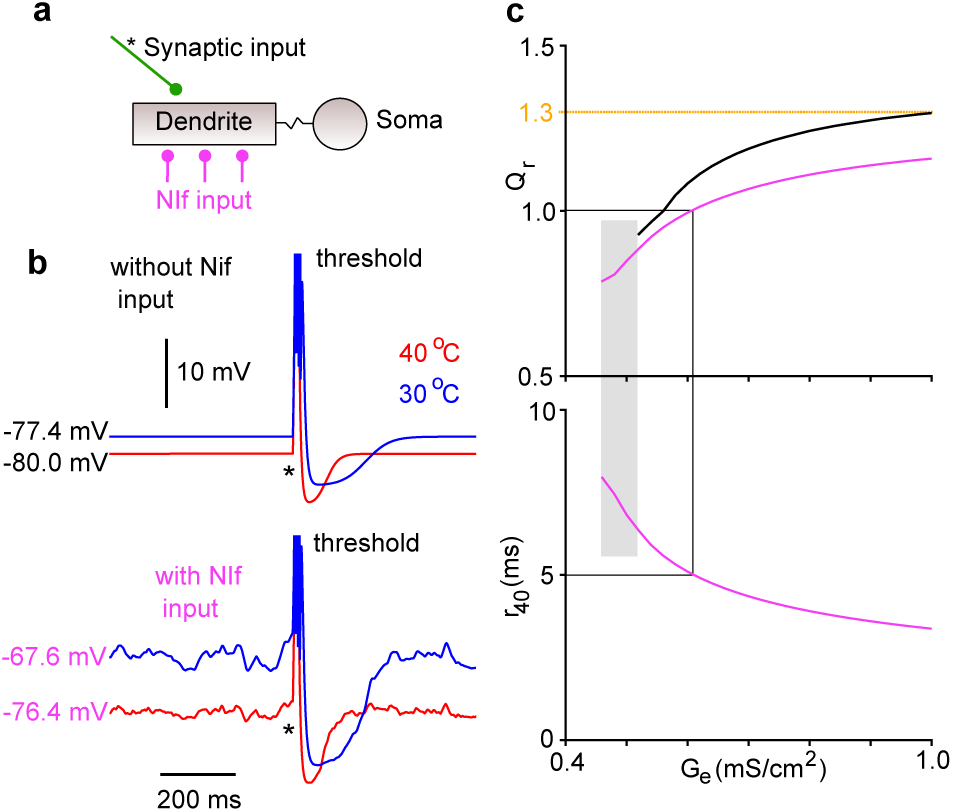
(a) The dendrite of the HVC_RA_ neuron receives tonic NIf input, which elevates the subthreshold membrane potential. (b) Membrane potential of the soma, with and without NIf input, at 40 °C and 30 °C. Cooling elevates the membrane potential more in the presence of NIf input than in its absence. Additionally, NIf input introduces membrane potential fluctuations. (c) With NIf input, the Q_r_ of the rise time is lower than without it, as observed in Q_r_ as a function of *G_e_*. NIf input also facilitates burst generation at lower *G_e_* values (shaded region). The bottom panel shows the rise time *r*_40_ in the presence of NIf input.

Cooling reduced the NIf synaptic conductance by a factor of Q_10_ = 1.3, but the frequency remained unchanged because NIf activity is not affected by HVC cooling. Additionally, the synaptic decay time constant increased by Q_10_ = 3. Without NIf inputs, cooling to 30 °C elevated the membrane potential by 2.6 mV (Fig. 4b). In contrast, with NIf input, the membrane potential was elevated by 8.8 mV (Fig. 4b). This indicates that NIf input enhances HVC_RA_ neuron excitability when cooled, thereby reducing the temperature sensitivity of the neuronal response. Indeed, Q_r_ is lower with NIf input compared to the case without it (Fig. 4c). Furthermore, NIf input enables the generation of burst responses at 40 °C even when *G_e_*is insufficient to induce bursts in its absence (shaded region in Fig. 4c). In this regime, the *r*_40_ of our model HVC_RA_ neurons was within the 5–10 ms range, and the synaptic response exhibited high temperature robustness (*Q_r_ <* 1).

Cooling enhances the EPSPs of NIf input for the same reason described in the previous section. This enhancement further increases the temperature robustness of the synaptic responses of HVC_RA_ neurons.

### Temperature robustness of the HVC synaptic chain network

To evaluate the temperature robustness of synaptic chain network dynamics, we compared the burst propagation time at 40 °C and 30 °C (Fig. 5a). Bursts were initiated at the head of the chain network and propagated along the synaptic connections between HVC_RA_ neurons. HVC_INT_ neurons provided feedback inhibition to regulate the dynamics. Burst propagation time along the network was measured as the difference in the median burst times between two groups of neurons along the chain (Fig. 5a).

**Figure 5:**
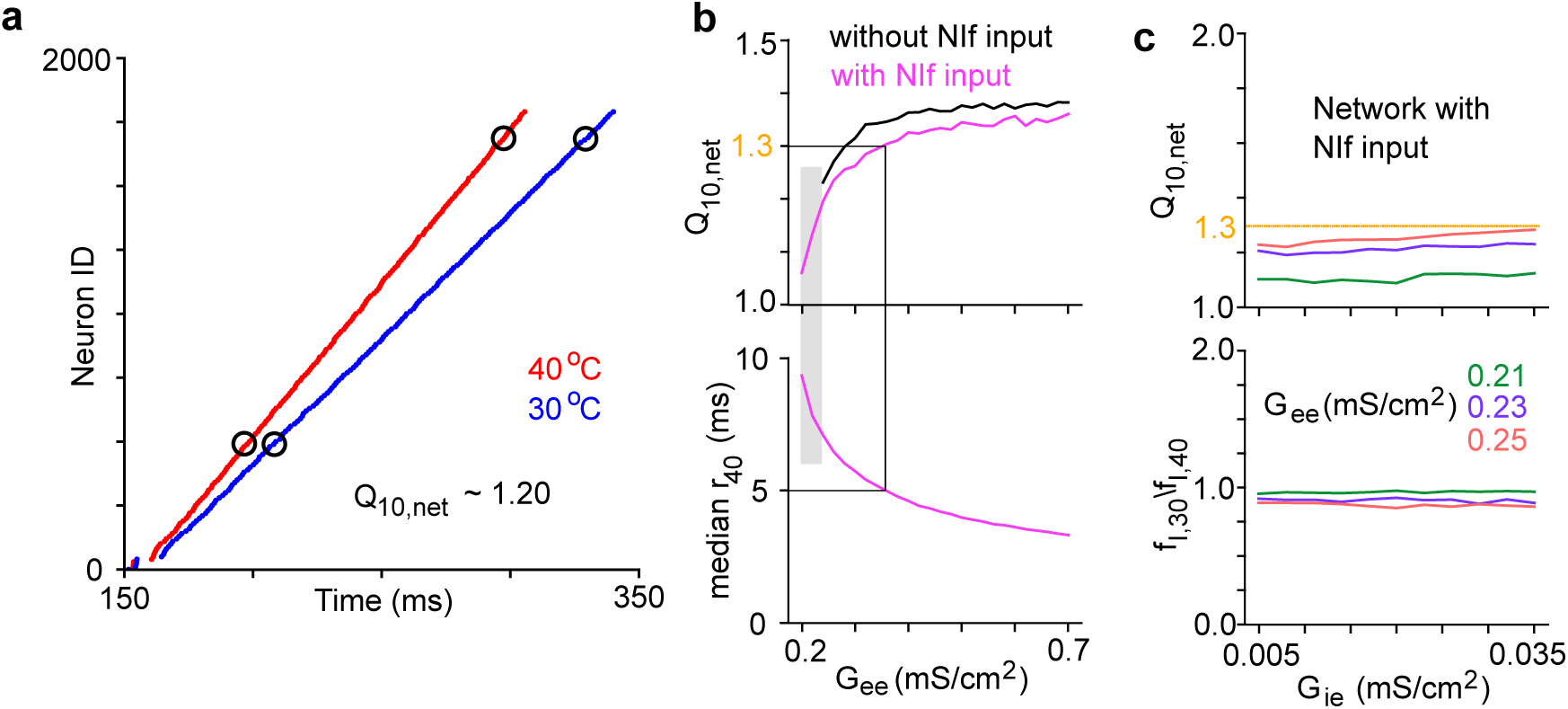
Temperature robustness of burst propagation for the synaptic chain network. (a) Raster plot of burst onset times for HVC_RA_ neurons at temperatures of 40 °C and 30 °C. The time required for bursts to propagate between two groups of HVC_RA_ neurons in the network (circled) was tracked. Upon cooling, the burst propagation time increased by a factor of Q_10,net_ = 1.2. (b) Q_10,net_ as a function of the connection strength *G_ee_* between HVC_RA_ neurons. With NIf input, Q_10,net_ is lower than without it. Below, the median membrane rise time *r*_40_ for 200 HVC_RA_ neurons with NIf input is shown. In the shaded region, NIf input is necessary for burst propagation at 40 °C. (c) Q_10,net_ for three excitation strengths with varying inhibition strength *G_ie_*. The *G_ee_* values lie within the shaded regime. Below, the ratios of the firing rates of HVC_INT_ neurons at 30 °C and 40 °C are shown.

As expected from our analysis above, the network dynamics slowed down with cooling but remained temperature-robust. The Q_10,net_ of the propagation time was approximately 1.2 (Fig. 5a), similar to the Q_10_ factor by which song tempo slowed in zebra finches during HVC cooling (Long and Fee, 2008). Therefore, our network simulation demonstrates that the Q_10_ factor of song tempo observed in zebra finches is consistent with the Q_10_ factor for burst propagation along a synaptic chain network localized within HVC.

We also investigated how Q_10,net_ depends on the connection strength *G_ee_*between HVC_RA_ neurons and NIf inputs (Fig. 5b). Similar to the case of a single HVC_RA_ neuron, increasing *G_ee_* led to an increase in Q_10,net_. In the absence of NIf input, Q_10,net_ increased from 1.23 and plateaued at 1.38. With NIf input, Q_10,net_ was lower, and increased from 1.06 and plateaued at 1.36. For values of *G_ee_* where the membrane rise time *r*_40_ at 40 °C was in the 5 - 10 ms range, Q_10,net_ was in the range 1.23 - 1.35 without NIf input and in the range 1.06 - 1.3 with NIf input. We next investigated the role of feedback inhibition on Q_10,net_ (Fig. 5c). In our model, HVC_INT_ and HVC_RA_ neurons were wired to ensure that inhibition exhibited gaps in the 20 ms window before HVC_RA_ neuron bursts (Fig. S2), consistent with experimental observations (Kosche et al., 2015). Thus, we expected that varying the inhibition strength *G_ie_* onto HVC_RA_ neurons would have minimal impact on Q_10,net_. We set the excitatory connections such that burst propagation depended on NIf input. Indeed, modifying *G_ie_* had little effect on Q_10,net_ (Fig. 5c). We also computed the ratio of the average firing rates of HVC_INT_ neurons, *f*_30_*/f*_40_, at 30 °C and 40 °C, finding it to be slightly below 1 (Fig. 5c). Cooling had little effect on the population firing rate of the HVC_INT_ neurons.

We have used Q_10_ = 3 for ion channel and synaptic dynamics so far. To assess the impact of this choice, we also examined the network dynamics by setting Q_10_ = 2, which is often used in modeling temperature effects (Hamaguchi et al., 2016). We found that the network dynamics became even more robust against cooling, with Q_10,net_ remaining below 1.3 across the entire range of *G_ee_* tested (Fig. S3).

## Discussion

Neuronal processes exhibit varying levels of temperature sensitivity. Cooling markedly slows ion channel dynamics (Schwarz and Eikhof, 1987; Sterratt, 2015; Pahlavan et al., 2023) and prolongs synaptic current decay (Q_10_ > 2) (Hestrin et al., 1990). In contrast, cooling has a moderate effect on ion channel conductances and spike propagation speed along axons (Q_10_ ∼ 1.3) (Hodgkin et al., 1952; Swadlow et al., 1981). Cooling the HVC region in songbirds moderately slows song speed (Q_10_ ∼ 1.3) (Long et al., 2010; Aronov and Fee, 2012; Zhang et al., 2017). We find that this temperature robustness of song speed is consistent with a model in which the song tempo is controlled by a synaptic chain network localized within the HVC (Jin et al., 2007; Long et al., 2010; Egger et al., 2020; Tupikov and Jin, 2021). This robustness arises from the reliance on axonal delays and the enhanced efficacy of EPSPs upon cooling.

Axonal delays between HVC_RA_ neurons are distributed over a range of 1 to 7.5 ms, with a mean of 3.3 ms (Egger et al., 2020). The synaptic chain network in our work follows the polychronous principle, where the postsynaptic neuron receives synchronous inputs from presynaptic neurons despite the distributed axonal delays (Izhikevich, 2006; Egger et al., 2020). Burst propagation time along the chain consists of two components: axonal delays from presynaptic to postsynaptic neurons and the neuronal integration time in postsynaptic neurons (Fig. 2e). Intra-cellular recordings of HVC_RA_ neurons in singing zebra finches have shown that their membrane potential rises from baseline over an interval of 5 - 10 ms before burst onset. Thus, the time spent in spike propagation along axons is comparable to the time spent in neuronal integration.

This reliance on axonal delays enables the network to leverage the temperature robustness of axonal conductance. In principle, local neural circuits can mitigate the impact of temperature fluctuations by relying more on axonal delays for computation.

Ion channel and synaptic decay dynamics rely on protein conformation changes and hence are highly temperature-sensitive (Sterratt, 2015). Indeed, the rates of any chemical reactions that depend on overcoming energy barriers follow the Arrhenius equation and depend exponentially on temperature (Liang, 2022). However, it is often incorrect to conclude that neural circuit dynamics and the behaviors encoded by such circuits must also be highly sensitive to temperature changes. Examples are abundant. The circadian clocks in mammalian central and peripheral tissues are almost perfectly temperature-compensated, with Q_10_ ∼ 1 (Reyes et al., 2008). The firing rate of grasshopper auditory receptor neurons is temperature robust, with Q_10_ ∼ 1.5 (Roemschied et al., 2014). In the pyloric circuit of the crab stomatogastric ganglion, the fraction of time the oscillator spends in bursting remains essentially constant (Q_10_ ∼ 1) over a wide range of temperature (Alonso and Marder, 2020). Spiking of hair cells in bullfrogs maintains sensitivity and precision to sounds over a wide range of temperature (Chen and Von Gersdorff, 2019). The wingbeat frequency of locusts changes only slightly with temperature (Q_10_ ∼ 1.15) (Robertson and Money, 2012).

Temperature-sensitive neuronal processes can exhibit counteracting properties that promote temperature robustness at the circuit level (Schapiro and Marder, 2024). In our case, cooling slows the closing of synaptic receptors, thereby enhancing synaptic transmission. Additionally, cooling elevates the resting membrane potential of neurons and reduces leak conductance, which enhances neuronal excitability, as observed in intracellular recordings of layer 2/3 pyramidal neurons in rat visual cortical slices (Volgushev et al., 2000). These effects counteract the slowing of neural integration processes and promote the temperature robustness of circuit dynamics. NIf has been shown to provide excitatory input to HVC that promote HVC activity during singing in zebra finches (Otchy et al., 2015). Our model demonstrates that this excitatory external input also enhances the robustness of synaptic chain dynamics against cooling.

Following experimental observations (Kosche et al., 2015), we wired the HVC_RA_ and HVC_INT_ neurons such that HVC_RA_ neurons receive less inhibition in the 20 ms window before the time of their bursts. Such gaps in inhibition prevent the feedback inhibition from interfering with burst generation in HVC_RA_ neurons (Kosche et al., 2015). In our case, these inhibition gaps ensure that the enhancement of inhibitory synapses upon cooling does not reduce temperature robustness of burst propagation. Indeed, if we randomly wire the HVC_RA_ and HVC_INT_ neurons, as done in previous models (Jin, 2009; Long et al., 2010; Egger et al., 2020), inhibition gaps do not emerge. In this scenario, we found that cooling increases the firing rate of HVC_INT_ neurons due to the enhanced efficacy of the excitatory synapses from HVC_RA_ neurons (Fig. S2d,e). Additionally, cooling enhances the efficacy of inhibitory synapses onto HVC_RA_ neurons. Together, these effects slow synaptic integration time and lead to an increase in Q_10,net_ (Fig. S2c). Thus, inhibition gaps play a crucial role in promoting the temperature robustness of the synaptic chain network. Such gaps can naturally arise through the self-organized process of wiring the synaptic chain network via the recruitment of newborn neurons that rely on their spontaneous activity (Tupikov and Jin, 2021).

Inhibition in HVC has been proposed to play an important role in models of generating variable syllable sequences in the songs of species such as the Bengalese finch (Jin, 2009). During the silent gap leading to probabilistic syllable transitions, inhibition facilitates a winner-take-all mechanism for selecting the next syllable among possible candidates (Jin, 2009). Upon cooling, enhanced inhibition could slow the selection dynamics, thereby prolonging the silent gap. Indeed, experiments that cooled the Bengalese finch HVC during singing observed a larger Q_10_ for silent gaps than for syllable durations (Zhang et al., 2017), in agreement with our analysis of the role of inhibition.

A previous study questioned the synaptic chain network in HVC as the mechanism for generating sparse burst sequences in HVC_RA_ neurons (Hamaguchi et al., 2016). The argument was that cooling HVC should result in dynamics with Q_10_ ∼ 2, since most biological processes have Q_10_ ∼ 2. This would contradict the observation that song Q_10_ is approximately 1.3.

The authors further proposed that HVC is part of a recurrent loop network involving HVC, RA, the brainstem, and the thalamic nucleus Uvaeformis (Uva), which projects back to HVC (Hamaguchi et al., 2016). Within this framework, bursts of HVC_RA_ neurons propagate through the loop, sequentially triggering HVC_RA_ bursts at subsequent moments.

While this argument is plausible, it overlooks the possibility that temperature-sensitive elements within neural circuits could counterbalance each other, rendering the circuit temperature-robust (Schapiro and Marder, 2024). Additionally, it does not account for the use of temperature-robust circuit elements such as axonal delays (Egger et al., 2020). Indeed, the temperature of the entire brain of a zebra finch can increase by up to 4 °C when a female is present (Aronov and Fee, 2012). The song tempo is faster, and the Q_10_ remains around 1.3, similar to the effect of cooling HVC alone (Aronov and Fee, 2012). This observation directly contradicts the intuitive expectation that Q_10_ ∼ 2.

The recurrent loop model predicts a prominent 50 Hz oscillation in the synaptic activity of HVC_RA_ neurons due to the time required for spikes to complete a round trip back to HVC through the loop (Hamaguchi et al., 2016). However, recordings of HVC_RA_ neuron activity in singing zebra finches have shown no such oscillations (Egger et al., 2020).

The ion channel Q_10_ can be temperature-dependent, often decreasing at higher temperatures (Pahlavan et al., 2023). Given that birds maintain a higher average body temperature (∼40 °C) compared to mammals (∼37 °C), it is plausible that the Q_10_ for ion channels in birds may be lower than in mammals. This, in turn, would reduce Q_10,net_, as shown in Fig. S3. Further experiments would be valuable in investigating this possibility.

In conclusion, we have identified neuronal mechanisms that support temperature-robust burst propagation along the synaptic chain network localized within HVC. This robustness, quantified by a Q_10_ for burst propagation, is in close agreement with the observed Q_10_ ∼1.3 for the song tempo. Consequently, our findings lend further support to the role of HVC as a critical brain nucleus involved in encoding the timing and moment-to-moment features of zebra finch song.

## Methods

### HVC neuron model

The computational models of HVC_RA_ and HVC_INT_ neurons follow previous works (Jin, 2009; Wittenbach et al., 2015; Tupikov and Jin, 2021), and details can be found therein. We further introduced an additional ion channel in the dendritic compartment of the HVC_RA_ neuron: the big-conductance (BK) calcium-dependent potassium channel. This channel was incorporated to ensure that calcium spikes in the dendrite produce somatic bursts of 4 - 5 sodium spikes at both 40 °C and 30 °C. The BK channel model follows the formulation presented in (Womack and Khodakhah, 2002). The current is given by

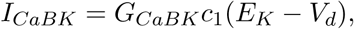

where *G_CaBK_* = 100 mS/cm^2^ is the channel conductance, *E_K_*= −90 mV is the reversal potential of the potassium channel, and *V_d_* is the dendritic membrane potential. The activation variable *c*_1_ follows

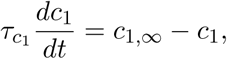

where the time constant is *τ*_*C*1_ ms, and the steady-state value is

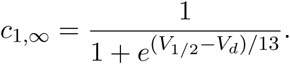

The half-activation voltage depends on the calcium concentration [*Ca*] and is given by

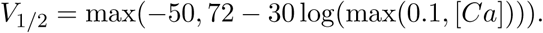

The calcium concentration [*Ca*] evolves according to

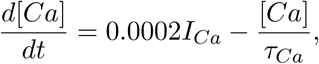

where *I_Ca_*is the calcium current, and *τ_Ca_*= 50 ms is the buffering time constant (Jin, 2009; Wittenbach et al., 2015; Tupikov and Jin, 2021).

### Synaptic dynamics

We used the kick-and-decay model for synaptic dynamics, similar to previous works (Jin, 2009; Wittenbach et al., 2015; Tupikov and Jin, 2021). When a spike arrives, the synaptic conductance *g* increases by a fixed value *G*, which represents the synaptic strength. Between spikes, *g* decays with a synaptic time constant *τ* . We set *τ* = 2 ms at 40 °C. With *Q* = 3 for *τ*, it increases to 6 ms at 30 °C.

### Noise model

Noisy fluctuations in neuronal membrane potentials were generated by random spike inputs to the compartments (Jin, 2009). For HVC_RA_ neurons, the dendritic and somatic compartments receive random spikes at a rate of 200 Hz, with conductance sampled uniformly from the interval (0, 0.06) mS/cm^2^. For HVC_INT_ neurons, the spike frequency was 300 Hz, with conductance sampled from the interval (0, 0.45) mS/cm^2^. With equal probability, these random spikes contributed to either excitatory or inhibitory conductances of the compartment. The model HVC_RA_ neuron’s soma exhibits subthreshold fluctuations on the order of ∼3 mV (Long et al., 2010; Mooney, 2000), while the model HVC_INT_ neuron exhibits spontaneous firing rates close to ∼10 Hz (Kozhevnikov and Fee, 2007).

### NIf input

NIf input was modeled as random excitatory spikes generated by a Poisson process with a frequency of 1500 Hz and a conductance uniformly sampled in the range (0, 0.005) *mS/cm*^2^. The spikes arrive at the dendritic compartment of HVC_RA_ neurons.

### Algorithm for wiring a polychronous synaptic chain network

Axonal delays of connections between HVC_RA_ neurons are distributed in the range of 1 - 7.5 ms (Egger et al., 2020). We wired the synaptic chain network to ensure that the postsynaptic neuron receives nearly synchronous inputs from presynaptic neurons, despite the distributed axonal delays (Egger et al., 2020). We used a method that is significantly simpler than the one proposed in previous work (Egger et al., 2020).

We wired *N* = 2000 HVC_RA_ neurons. We created 50 bins, each with a duration of 1 ms, covering a total duration of 50 ms. This represents the overall contribution of axonal delays to the burst propagation time in our network. We assigned N_bin_ = 40 neurons to each bin, so that the neurons in the *i*-th bin had putative burst onset time in the interval *t* = (*i, i* + 1) ms. Each neuron sent out N_out_ = 50 connections. The neurons in the first bin were designated as starter neurons. These neurons were activated by external inputs to initiate burst propagation in the chain network. The neurons in the last four bins were designated as terminating neurons and do not send out connections.

We iterated over all non-terminating neurons in the order of their assigned putative burst onset times. A neuron to be wired was called a source neuron, and its putative burst onset time was denoted as *t*_source_. To establish a connection, we sampled the delay time *d* from a log-normal distribution with a mean of 3 ms and a standard deviation of 1.5 ms. We then set the target putative burst onset time as *t*_target_ = *t*_source_ + *d.* Next, we randomly selected a neuron whose putative burst onset time fell within the range *t*_target_ ± *δ* and established a connection from the source neuron to this target neuron with an axonal delay of *d*, where *δ* = 0.25 ms is the tolerance.

This procedure was repeated until each source neuron formed all N_out_ connections. Although there was no explicit constraint on the number of input connections a neuron could receive, the number of input connections per neuron was approximately N_out_, except for neurons in the first few bins due to the limited availability of suitable source neurons.

For each neuron, the total excitatory conductance was set to *G_ee_*. The synaptic conductance of each incoming connection was then set to *G_ee_/n_in_*, where *n_in_* is the number of incoming connections. This ensured that the synaptic integration time for neurons remained roughly the same, despite fluctuations in *n_in_*. Additionally, the synchrony of presynaptic spike arrival was preserved, even though our wiring process did not explicitly take synaptic integration time into account.

### Wiring inhibitory neurons to the synaptic chain network

In previous studies, HVC_INT_ neurons were randomly connected to HVC_RA_ neurons (Jin, 2009; Long et al., 2010). This resulted in random inhibition, where the timing of feedback inhibition was uncorrelated with the burst times of HVC_RA_ neurons. However, experimental observations indicate that inhibition is absent before the onset of bursts in HVC_RA_ neurons (Kosche et al., 2015). Moreover, random inhibition led to elevated membrane potentials in HVC_INT_ neurons throughout burst propagation. In contrast, experimental data showed that membrane potentials were elevated only during several short segments, corresponding to the periods when HVC_INT_ neurons spiked. To align our model with experimental findings, we structured the connections of HVC_INT_ neurons to generate gapped inhibition.

Our network contained 550 HVC_INT_ neurons. Before establishing the HVC_RA_ -to-HVC_INT_ connections, we determined the burst onset times of HVC_RA_ neurons by simulating the dynamics of the synaptic chain network with *G_ee_* = 0.25 mS/cm^2^. To wire an HVC_RA_ neuron, we randomly selected 3 – 5 segments within the burst propagation period, with each segment lasting 1 ms (Fig. S2b). HVC_RA_ neurons whose burst onset times fell within these segments were connected to the HVC_INT_ neuron. Axonal delays were sampled from a log-normal distribution with a mean of 1.55 ms and a standard deviation of 0.6 ms (Fig. S2h), while connection strengths were uniformly selected from the interval (0*, G_ei_*). These segments defined the firing times of the HVC_INT_ neuron, with variations across runs due to noise. This process was repeated for each HVC_INT_ neuron.

We then established the HVC_INT_ -to-HVC_RA_ connections. For each HVC_RA_ neuron, we excluded HVC_INT_ neurons whose spike times occurred within the 20 ms window preceding the burst onset of the HVC_RA_ neuron. From the remaining HVC_INT_ neurons, we randomly selected connections with a probability of *p* = 0.1. Axonal delays were sampled from a log-normal distribution with a mean of 1.1 ms and a standard deviation of 0.5 ms (Fig. S2i), while connection strengths were uniformly sampled from the interval (0*, G_ie_*). This process was repeated for all HVC_RA_ neurons.

For comparison with gapped inhibition, we also wired the HVC_INT_ and HVC_RA_ neurons randomly. The probability of a connection from an HVC_RA_ neuron to an HVC_INT_ neuron was set to 0.03, while the probability of a connection from an HVC_INT_ neuron to an HVC_RA_ neuron was set to 0.045. The number of input connections determined by these probabilities matched that of the network wired with gapped inhibition. To ensure that the average firing rate of HVC_INT_ neurons was roughly equivalent to that in the gapped inhibition wiring, we set *G_ei_* = 0.35 mS/cm^2^.

### Synaptic response of an HVC_RA_ neuron

In the analysis of the synaptic response of an HVC_RA_ neuron, an excitatory spike was delivered at 150 ms to the dendritic compartment, with no noise spikes present. The membrane potential rise time, *r_T_*, at temperature *T* was defined as the duration from the spike input to the peak membrane potential (or the time until the membrane potential crossed -20 mV if the soma produced a burst of spikes). The input strength, *G_e_*, was varied within the range of 0–5 mS/cm^2^. The Q_10_ of the rise time was defined as Q_r_ = *r*_30_*/r*_40_.

### Synaptic response of an HVC_RA_ neuron with NIf input

The influence of NIf on Q_r_ was studied in the regime where the HVC_RA_ neuron responded with a somatic burst of spikes at both 40 °C and 30 °C. NIf input was modeled as random excitatory spikes with a frequency of 1500 Hz using a Poisson process. The input strength of each spike was selected uniformly from the interval (0, 0.005) mS/cm^2^. Noisy spikes were present. The strength of the synaptic input, *G_e_*, varied from 0.4 to 1 mS/cm^2^. The results were obtained by averaging 50 runs of the dynamics.

### Calculating Q_10,net_

The Q_10_ factor of burst propagation time was defined as Q_10,net_ = Δ*t*_30_*/*Δ*t*_40_, where Δ*t*_30_ and Δ*t*_40_ represent the differences in the median burst times of two groups of 100 HVC_RA_ neurons at 30 °C and 40 °C, respectively. Network activity was initiated by applying a strong synaptic input of strength 1.0 mS/cm^2^ to the starter neurons at *t* = 150 ms.

The dependence of Q_10,net_ on the excitatory connection strength, *G_ee_*, between HVC_RA_ neurons was studied by varying *G_ee_*from 0.2 to 0.7 mS/cm^2^. The connection strength *G_ei_*, from HVC_RA_ to HVC_INT_ neurons, was set to 0.15 mS/cm^2^, while the connection strength *G_ie_*, from HVC_INT_ to HVC_RA_ neurons, was set to 0.023 mS/cm^2^.

The role of NIf input was studied by applying random NIf spikes to the dendritic compartment of HVC_RA_ neurons in the same manner as in the single-neuron case described above.

The population firing rates, *f_I,T_*, of HVC_INT_ neurons were computed by calculating the average spike rates of HVC_INT_ neurons within the intervals Δ*t_T_*of the two HVC_RA_ neuron groups.

### Analytical analysis of the leaky integration dynamics

The equation for membrane potential is

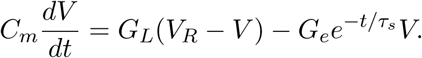

Here *C_m_*is the membrane capacitance, *G_L_*is the leak conductance, *V_R_*is the resting membrane potential, *G_e_* is the synaptic conductance, and *τ_s_* is the synaptic decay time constant.

It is convenient to make this equation dimensionless by defining a few quantities:

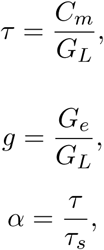

and

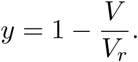

Scale time as

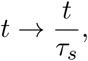

then the equation becomes

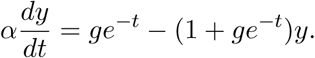

In the *t* − *y* plane, the nullcline is given by

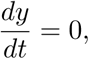

or

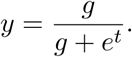

This is where the membrane potential turns around. On the left of the nullcline, *y*(*t*) increases with time. On the right of the nullcline, *y*(*t*) decreases with time. The nullcline is a monotonically decreasing function of *t*.

Because of the uniqueness theorem, solutions of the first order differential equations with the same initial condition cannot cross each other. At *t* = 0, we have

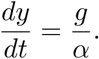

Therefore, small *α* results in a larger derivative. This makes the EPSP of a small *α* entirely above that of large *α*. In other words, small *α* means larger EPSP. Since cooling keeps *g* unchanged but decreases *α*, it results in a larger EPSP.

## Conflict of Interest

The authors declare no competing financial interests.

## Acknowledgements

Supported by NSF Award EF-1822476 (DZJ). The funders had no role in study design, data collection and analysis, decision to publish, or preparation of the manuscript. We thank Michael Long for useful discussions and suggestions.

---

**Figure S1:**
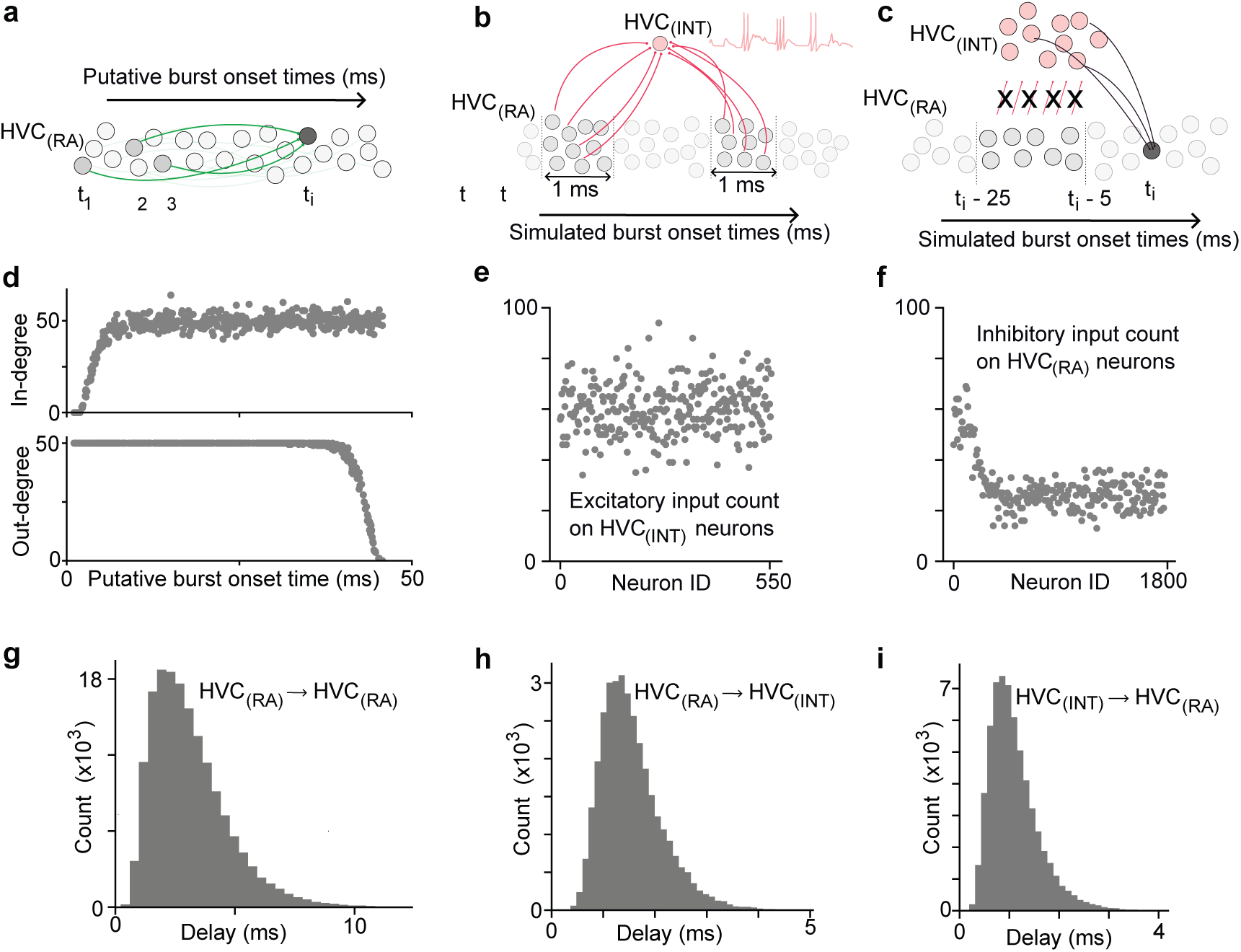
Wiring the synaptic chain network. (a) HVC_RA_ neurons are wired into a polychronous network. Each neuron is assigned a putative burst onset time, and presynaptic neurons are selected to ensure that burst spikes arrive at the postsynaptic neuron synchronously, despite axonal delays. (b) HVC_INT_ neurons are driven by multiple groups of HVC_RA_ neurons, with each group firing within a 1 ms window. (c) HVC_INT_ neurons provide inhibitory input to HVC_RA_ neurons but are wired in such a way that inhibition gaps occur just before HVC_RA_ burst onsets. (d) Number of incoming and outgoing connections for HVC_RA_ neurons in the network. (e, f) Number of excitatory inputs received by HVC_INT_ neurons and the number of inhibitory inputs received by HVC_RA_ neurons, respectively. (g, h, i) Distribution of axonal delays used in the wired network.

**Figure S2:**
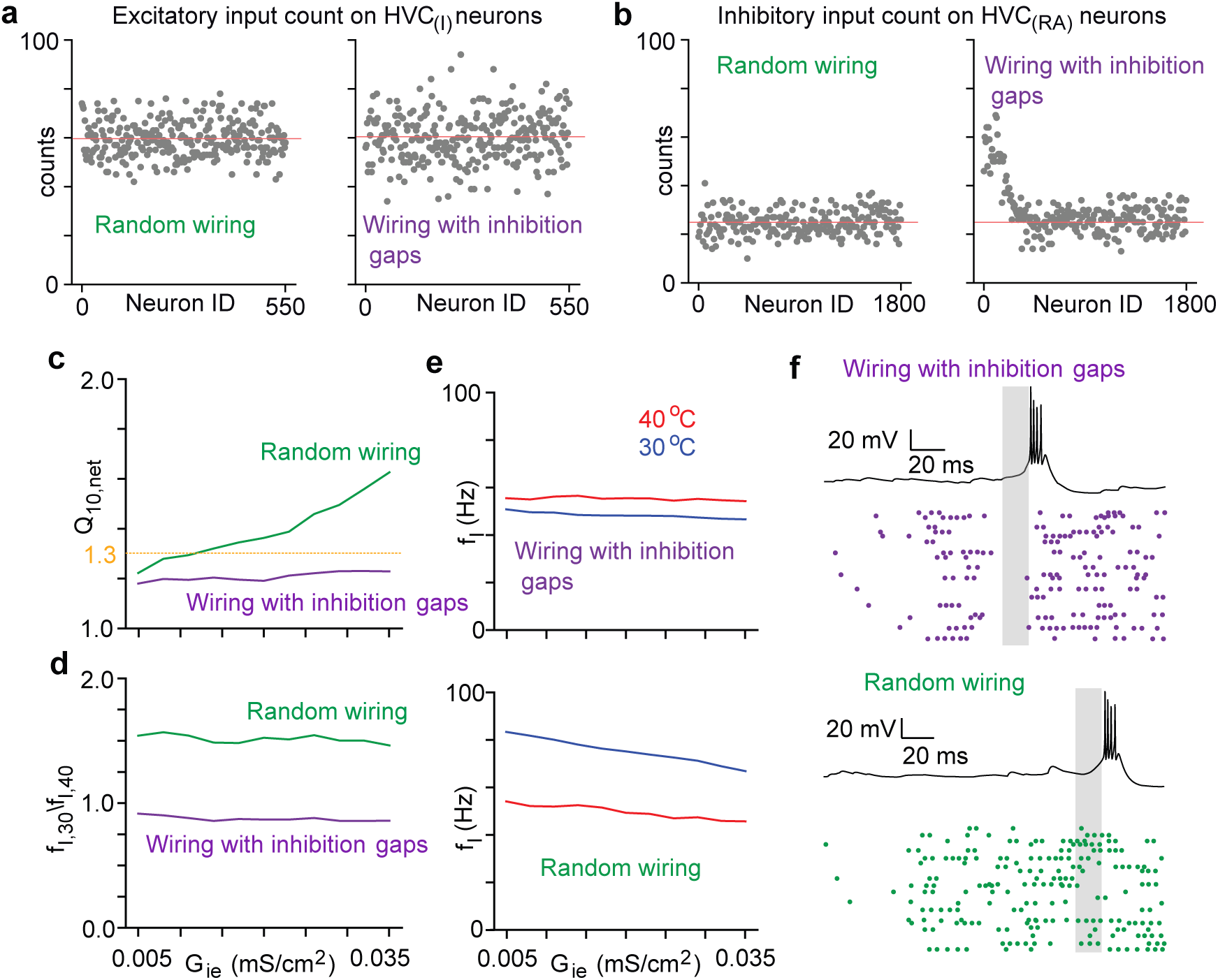
Comparing gapped inhibition with global inhibition. (a, b) The networks were structured such that the number of connections an HVC_INT_ neuron receives from HVC_RA_ neurons and the number of connections it makes onto HVC_RA_ neurons are similar. (c) Q_10,net_ as a function of *G_ie_* for the two networks. Purple: gapped inhibition; green: random inhibition. (d) Ratios of the firing rates of HVC_INT_ neurons, *f*_30_*/f*_40_, for the two networks. The firing rate increases with cooling in the random inhibition case (green) but remains unchanged in the gapped inhibition case (purple). (e) Firing rates of HVC_INT_ neurons in the two networks. (f) Example membrane potential traces for an HVC_RA_ neuron’s soma, along with raster plots of HVC_INT_ neurons that provide inhibitory input to the HVC_RA_ neuron. In the gapped inhibition case, no HVC_INT_ neuron spikes occur immediately before an HVC_RA_ neuron bursts (gray region), whereas in the random inhibition case, such spikes are present.

**Figure S3:**
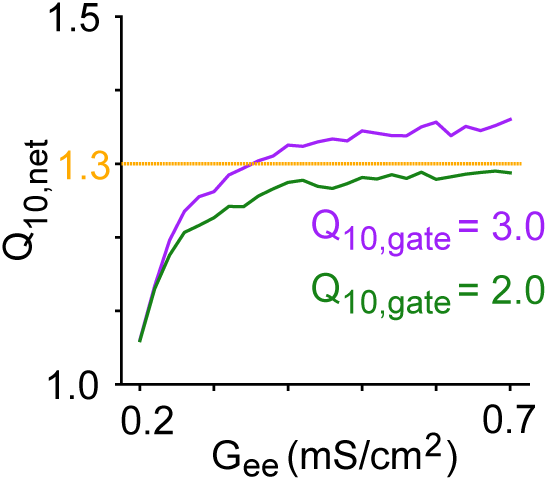
The effect of lowering Q_10_ for ion channel and synaptic dynamics to 2. The Q_10_ factor for the ion channel and synaptic dynamics, Q_10,gate_, was reduced from 3 to 2. The network Q_10,net_ is lower when Q_10,gate_ = 2 (green line) than when Q_10,gate_ = 3 (purple line) for the entire range of the connection strength, *G_ee_*, between the HVC_RA_ neurons.

## Notes

### Competing Interest Statement

The authors have declared no competing interest.

